# The Polycomb group protein Ring1 regulates dorsoventral patterning of the mouse telencephalon

**DOI:** 10.1101/639492

**Authors:** Hikaru Eto, Yusuke Kishi, Haruhiko Koseki, Yukiko Gotoh

**Author notes:** Correspondence (Y.K.).

## Abstract

Patterning of the dorsal-ventral (D-V) axis of the mammalian telencephalon is fundamental to the formation of distinct functional regions including the neocortex and ganglionic eminences. Morphogenetic signaling by bone morphogenetic protein (BMP), Wnt, Sonic hedgehog (Shh), and fibroblast growth factor (FGF) pathways determines regional identity along this axis. It has remained unclear, however, how region-specific expression patterns of these morphogens along the D-V axis are established, especially at the level of epigenetic (chromatin) regulation. Here we show that epigenetic regulation by Ring1, an essential Polycomb group (PcG) protein, plays a key role in formation of ventral identity in the mouse telencephalon. Deletion of the *Ring1b* or both *Ring1a* and *Ring1b* genes in neuroepithelial cells of the mouse embryo attenuated expression of the gene for Shh, a key morphogen for induction of ventral identity, and induced misexpression of dorsal marker genes including those for BMP and Wnt ligands in the ventral telencephalon. PcG protein–mediated trimethylation of histone H3 on lysine-27 (H3K27me3) was also apparent at BMP and Wnt ligand genes in wild-type embryos. Importantly, forced activation of Wnt or BMP signaling repressed the expression of *Shh* in organotypic and dissociated cultures of the early-stage telencephalon. Our results thus indicate that epigenetic regulation by PcG proteins—and, in particular, that by Ring1— confers a permissive state for the induction of *Shh* expression through suppression of BMP and Wnt signaling pathways, which in turn allows the development of ventral identity in the telencephalon.

## Introduction

In vertebrate embryos, the telencephalon is formed at the most anterior portion of the developing central nervous system (CNS). The cerebral cortex (CTX) and ganglionic eminences (GEs) are derived from the dorsal telencephalon (the pallium) and the ventral telencephalon (the subpallium), respectively (Campbell, 2003; Hebert and Fishell, 2008). Whereas neural stem-progenitor cells (NPCs) in the CTX produce excitatory cortical neurons, astrocytes, and some late-born oligodendrocytes, those in the GE produce local neurons and glial cells that constitute the basal ganglia as well as inhibitory neurons and early-born oligodendrocytes that migrate tangentially to the CTX. Regulation of dorsal-ventral (D-V) patterning is thus fundamental to development of the telencephalon.

The regional identity of telencephalic NPCs along the D-V axis is determined by region-specific transcription factors such as Pax6 and Emx1/2 in the CTX, Gsx2 in the lateral and medial GE (LGE and MGE, respectively), and Nkx2.1 in the MGE (Corbin et al., 2003; Kroll and O’Leary, 2005; Simeone et al., 1992; Sussel et al., 1999; Wigle and Eisenstat, 2008). Mutual gene repression by Pax6 and Gsx2 contributes to establishment of the D-V boundary (Corbin et al., 2003). *Neurog1* and *Neurog2*, proneural genes in the CTX, and *Ascl1*, a proneural (and oligodendrogenic) gene in the GE, are expressed according to the regional identity of NPCs in mouse embryos (Casarosa et al., 1999; Fode et al., 2000; Toresson et al., 2000).

Telencephalic regionalization along the D-V axis begins before closure of the neural tube, which occurs around embryonic day (E) 9.0 in mice, and is established before the onset of neurogenesis at ∼E10. The initial stages of D-V patterning are controlled by secreted morphogenetic signals (morphogens) that spread over various distances. The combination of the activities of different morphogens gives rise to distinct expression patterns of region-specific transcription factors in the telencephalon (Gupta and Sen, 2016; Harrison-Uy and Pleasure, 2012; Lupo et al., 2006; Sur and Rubenstein, 2005), as is also the case in the vertebrate spinal cord (Andrews et al., 2019) and in invertebrate embryos (Briscoe and Small, 2015). Bone morphogenetic proteins (BMPs), Wnt ligands, Sonic hedgehog (Shh), and fibroblast growth factor 8 (FGF8) are among the morphogens involved in D-V patterning in the mammalian telencephalon.

The dorsal midline regulates dorsal patterning of the telencephalon (Monuki et al., 2001). BMPs (BMP4, −5, −6, and −7) are secreted from the dorsal midline and paramedial neuroectoderm in the prospective forebrain (Furuta et al., 1997) and play pivotal roles in such patterning through induction of target genes including *Msx1*, *Lmx1a*, and *Wnt3a* (Cheng et al., 2006; Currle et al., 2005; Fernandes et al., 2007; Furuta et al., 1997; J. A. Golden et al., 1999; Hebert et al., 2002; Panchision et al., 2001). Knockout of BMP receptors thus results in loss of the dorsal-most structures of the telencephalon including the cortical hem and choroid plexus (Fernandes et al., 2007). Wnt ligands are also expressed in the dorsal region (Wnt1, −3, −3a, and −7b in the dorsal telencephalic roof plate at E9.5; Wnt2b, −3a, −5a, −7b, and 8b in the cortical hem and Wnt7a and −7b in the CTX at later stages) and contribute to aspects of dorsal patterning such as formation of the cortical hem and the CTX through induction of various transcription factors including Lef1 as well as Emx1/2, Pax6, and Gli3, respectively (Backman et al., 2005; Galceran et al., 2000; Harrison-Uy and Pleasure, 2012; Hasenpusch-Theil et al., 2012). In addition, Wnt signaling increases the activity of Lhx2, a selector gene for the CTX (Chou and Tole, 2019; Hsu et al., 2015). Suppression of the Wnt signaling pathway increases expression of ventral-specific genes throughout the dorsal pallium, indicating the importance of such signaling in dorsal patterning (Backman et al., 2005).

Shh, on the other hand, plays a major role in ventral patterning of the telencephalon (Blaess et al., 2014). Shh is secreted initially from the anterior mesendoderm or the prechordal plate (Aoto et al., 2009), then from the ventral hypothalamus, and finally from the rostroventral telencephalon including the preoptic area and MGE (Ericson et al., 1995; Fuccillo et al., 2004; Mathieu et al., 2002; Rohr et al., 2001; Shimamura et al., 1995). Shh signaling induces the expression of ventral transcription factors such as Foxa2, Nkx2.1, and Gsx2 and suppresses the repressor activity of Gli3, which is crucial for development of the dorsal telencephalon (Jeong and Epstein, 2003; Kuschel et al., 2003; Rallu et al., 2002; Shimamura and Rubenstein, 1997). FGF8 expressed in the anterior neural ridge also contributes to formation of the ventral telencephalon. FGF signaling is required for ventral expression of Shh and Nkx2.1 (Gutin et al., 2006; Shinya et al., 2001; Storm et al., 2006), and, conversely, Shh is required for maintenance of FGF8 expression at the anterior neural ridge (Hayhurst et al., 2008; Ohkubo et al., 2002; Rash and Grove, 2007).

Expression of BMP ligands and activation of BMP signaling are confined to the dorsal midline, with this confinement being critical for development of the ventral telencephalon, given that ectopic BMP signaling can suppress the expression of ventral morphogenetic factors such as Shh and FGF8 as well as that of the ventral transcription factor Nkx2.1 in the chick forebrain (J. A. Golden et al., 1999; Ohkubo et al., 2002). It is also important that Wnt signaling be confined to the dorsal pallium, given that ectopic activation of such signaling suppresses ventral specification in the developing mouse telencephalon (Backman et al., 2005). Mutual inhibition between dorsal and ventral morphogenetic factors explains in part the regional confinement of BMP and Wnt signaling (Huang et al., 2007; Storm et al., 2003). However, it has remained unclear whether any epigenetic factors (histone modifiers) participate in the establishment and maintenance of regional identity along the D-V axis of the developing telencephalon.

Polycomb group (PcG) proteins are repressive epigenetic factors that consist of two complexes, PRC1 and PRC2. These complexes catalyze the ubiquitylation of histone H2A at lysine-119 (H2AK119ub) and the trimethylation of histone H3 at lysine-27 (H3K27me3), respectively (Di Croce and Helin, 2013; Simon and Kingston, 2013). PcG proteins were first identified as transcriptional repressors of *Hox* genes in *Drosophila melanogaster*. These genes maintain regional identity along the anterior-posterior (A-P) axis in the fly embryo (Maeda and Karch, 2009). In mammals, PcG proteins also contribute to maintenance of the A-P axis during embryogenesis through repression of *Hox* genes in the neural tube (Chambeyron et al., 2005) as well as through that of forebrain-related genes in the midbrain (Zemke et al., 2015). Moreover, PcG proteins participate in cell subtype specification in the spinal cord (M. G. Golden and Dasen, 2012) and in the CTX in a manner dependent on temporal codes (Hirabayashi et al., 2009; Morimoto-Suzki et al., 2014; Pereira et al., 2010; Sparmann et al., 2013; Tsuboi et al., 2018). However, it has remained unknown whether PcG proteins regulate D-V patterning of the mammalian CNS including the telencephalon.

We now show that Ring1, an E3 ubiquitin ligase and essential component of PRC1 (de Napoles et al., 2004; Wang et al., 2004), is required for formation of the ventral telencephalon. Neural-specific ablation of Ring1B or of both Ring1A and Ring1B thus attenuated expression of ventral-specific genes such as *Gsx2*, *Nkx2.1*, and *Ascl1* as well as increased that of dorsal-specific genes such as *Pax6*, *Emx1*, and *Neurog1* in the telencephalon of mouse embryos. We found that *Shh* expression was markedly reduced, whereas BMP and Wnt signaling pathways were activated, in the ventral telencephalon of such Ring1B knockout (KO) or Ring1A/B double knockout (dKO) embryos. Moreover, Ring1B and H3K27me3 were found to be enriched at the promoters of several BMP and Wnt ligand genes in the telencephalon of wild-type (WT) embryos. Consistent with these results, forced activation of BMP or Wnt signaling suppressed *Shh* expression in explant cultures prepared from the embryonic telencephalon. Overall, our findings indicate that Ring1 establishes a permissive state for *Shh* expression in the ventral region of the telencephalon through suppression of BMP and Wnt signaling in this region.

## Results

### Deletion of *Ring1* in the neuroepithelium results in morphological defects in the telencephalon

To investigate the role of PcG proteins in the early stage of mouse telencephalic development, we deleted *Ring1b* with the use of the *Sox1-Cre* transgene, which confers expression of Cre recombinase in the neuroepithelium from before E8.5 (Takashima, 2007). We confirmed that expression of the Ring1B protein in the telencephalic wall was greatly reduced in *Ring1b*^flox/flox^;*Sox1-Cre* (Ring1B KO) mice at E10 compared with that in *Ring1b*^flox/flox^ or *Ring1b*^flox/+^ (control) mice, whereas the abundance of Ring1B in tissues outside of the telencephalic wall appeared unchanged in the Ring1B KO embryos (Figure S1A). The expression of Ring1B in the telencephalic wall was also greatly reduced in mice lacking both Ring1B and its homolog Ring1A (*Ring1a*^−/−^;*Ring1b*^flox/flox^;*Sox1-Cre*, or Ring1A/B dKO, mice) compared with that in *Ring1a*^−/−^;*Ring1b*^flox/flox^ or *Ring1a*^−/−^;*Ring1b*^flox/+^ (Ring1A KO) mice (Figure S1B). The level of H2AK119ub (a histone modification catalyzed by Ring1) in the telencephalic wall was reduced in Ring1B KO mice and, to a greater extent, in Ring1A/B dKO mice at E10 (Figure 1A–D). These results were consistent with the notion that Ring1A and Ring1B have overlapping roles in H2A ubiquitylation and that Ring1B makes a greater contribution to this modification than does Ring1A (Simon and Kingston, 2013).

**Figure 1.**
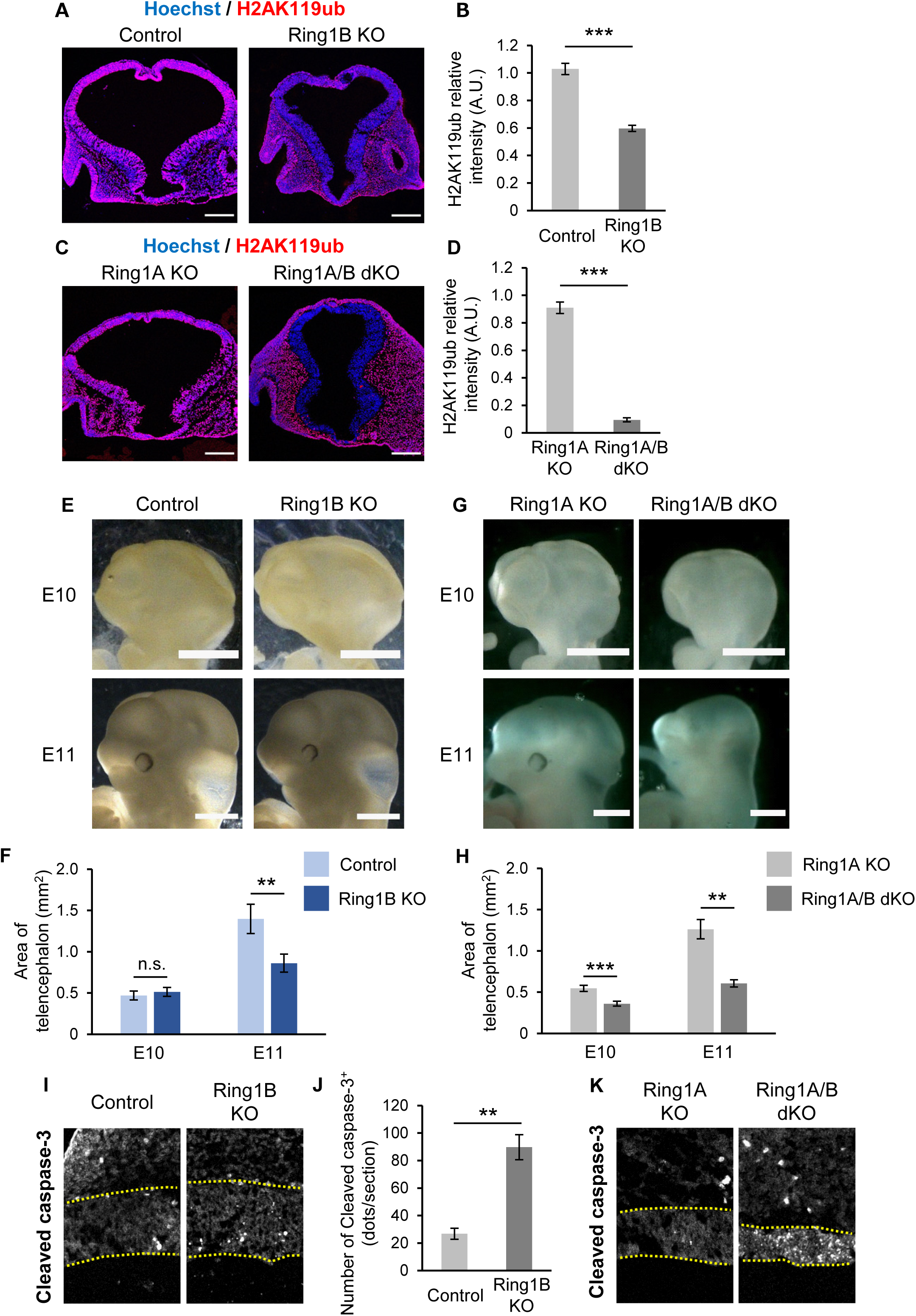
Deletion of *Ring1* in neural tissues results in morphological defects in the telencephalon. (A, C) Coronal sections of the brain of control (*Ring1b*^flox/flox^ or *Ring1b*^flox/+^) or Ring1B KO (*Ring1b*^flox/flox^;*Sox1-Cre*) mice (A) or of Ring1A KO (*Ring1a*^−/−^;*Ring1b*^flox/flox^ or *Ring1a*^−/−^;*Ring1b*^flox/+^) or Ring1A/B dKO (*Ring1a*^−.−^;*Ring1b* ^flox/flox^;*Sox1-Cre*) mice (C) at E10 were subjected to immunohistofluorescence analysis with antibodies to H2AK119ub. Nuclei were counterstained with Hoechst 33342. Scale bars, 200 µm. (B, D) The ratio of the average immunostaining intensity of H2AK119ub for the entire telencephalon to that for nonneural tissues adjacent to the ventral telencephalon was determined as relative intensity (A.U., arbitrary units) for images similar to those in (A) and (C). Multiple sections of the telencephalon along the rostrocaudal axis were examined for each embryo. Data are means ± s.e.m. of values for three embryos. ***P < 0.001 (two-tailed Student’s unpaired *t* test). (E, G) Images of the brain of control or Ring1B KO mice (E) or of Ring1A KO or Ring1A/B dKO mice (G) at E10 and E11. Scale bars, 1 mm. (F, H) Quantification of the lateral projected area of the telencephalon at E10 and E11 for images similar to those in (E) and (G). Data are means ± s.e.m. of averaged values from four litters (2-9 embryos per litter). **P < 0.01, ***P < 0.001; ns, not significant (two-tailed Student’s paired *t* test). (I, K) Coronal sections of the brain of control or Ring1B KO mice (I) or of Ring1A KO or Ring1A/B dKO mice (K) at E10 were subjected to immunohistofluorescence analysis with antibodies to the cleaved form of caspase-3. A region (200 by 300 µm) of the ventral telencephalon corresponding to the LGE is shown for each genotype. The neural tissue (the telencephalic wall) is demarcated by the yellow dotted lines. (J) The average number of cleaved caspase-3 signals in the control and Ring1B KO mouse telencephalon was determined with coronal section images. Data are means ± s.e.m. from three embryos. **P < 0.01 (two-tailed Student’s unpaired *t* test).

We found that *Ring1b* deletion with the use of *Sox1-Cre* resulted in a significant reduction in the size of the telencephalon at E11 (Figure 1E, F). Deletion of *Ring1b* at E13.5 with the use of the *Nestin-CreERT2* transgene was previously found not to substantially affect the morphology or size of the telencephalon (Hirabayashi et al., 2009), indicating that Ring1B functions during the early stage of telencephalon development. The reduction in telencephalon size was also pronounced in Ring1A/B dKO mice compared with Ring1A KO mice (Figure 1G, H). The number of cells positive for the cleaved form of caspase-3 (a marker for apoptosis) in the telencephalic wall was increased in Ring1B KO mice and, to a greater extent, in Ring1A/B dKO mice at E10 (Figure 1I–K), suggesting that Ring1 is necessary for the survival of telencephalic cells during the early stage of development and that the reduction in telencephalic size induced by *Ring1* deletion is due, at least in part, to the aberrant induction of apoptosis.

### Deletion of *Ring1* attenuates the expression of ventral-specific transcription factors in NPCs of the ventral telencephalon

We next investigated whether *Ring1* deletion might affect the D-V axis of the telencephalon by determining the expression of region-specific transcription factors. We examined embryos mostly at E10 given the apparently normal telencephalic size in Ring1B KO mice at this stage. Nkx2.1 is a transcription factor specifically expressed in the MGE (Sussel et al., 1999), and we found that the expression of this protein was greatly diminished in Ring1B KO mice at E10 (Figure 2A). The dorsal border of the Nkx2.1 expression domain was thus shifted ventrally and the abundance of Nkx2.1 within this domain was also reduced by *Ring1b* deletion (Figure 2B, C). The extent of the loss of Nkx2.1 expression appeared greater in Ring1A/B dKO mice than in Ring1B KO mice (Figure 2D–F). In addition to its effect on Nkx2.1 expression at the protein level, *Ring1b* deletion resulted in marked down-regulation of *Nkx2.1* mRNA in CD133^+^ NPCs isolated from the GE of the telencephalon at E11 (Figure 2I). These results thus indicated that Ring1 is necessary for ventral expression of the MGE marker Nkx2.1.

**Figure 2.**
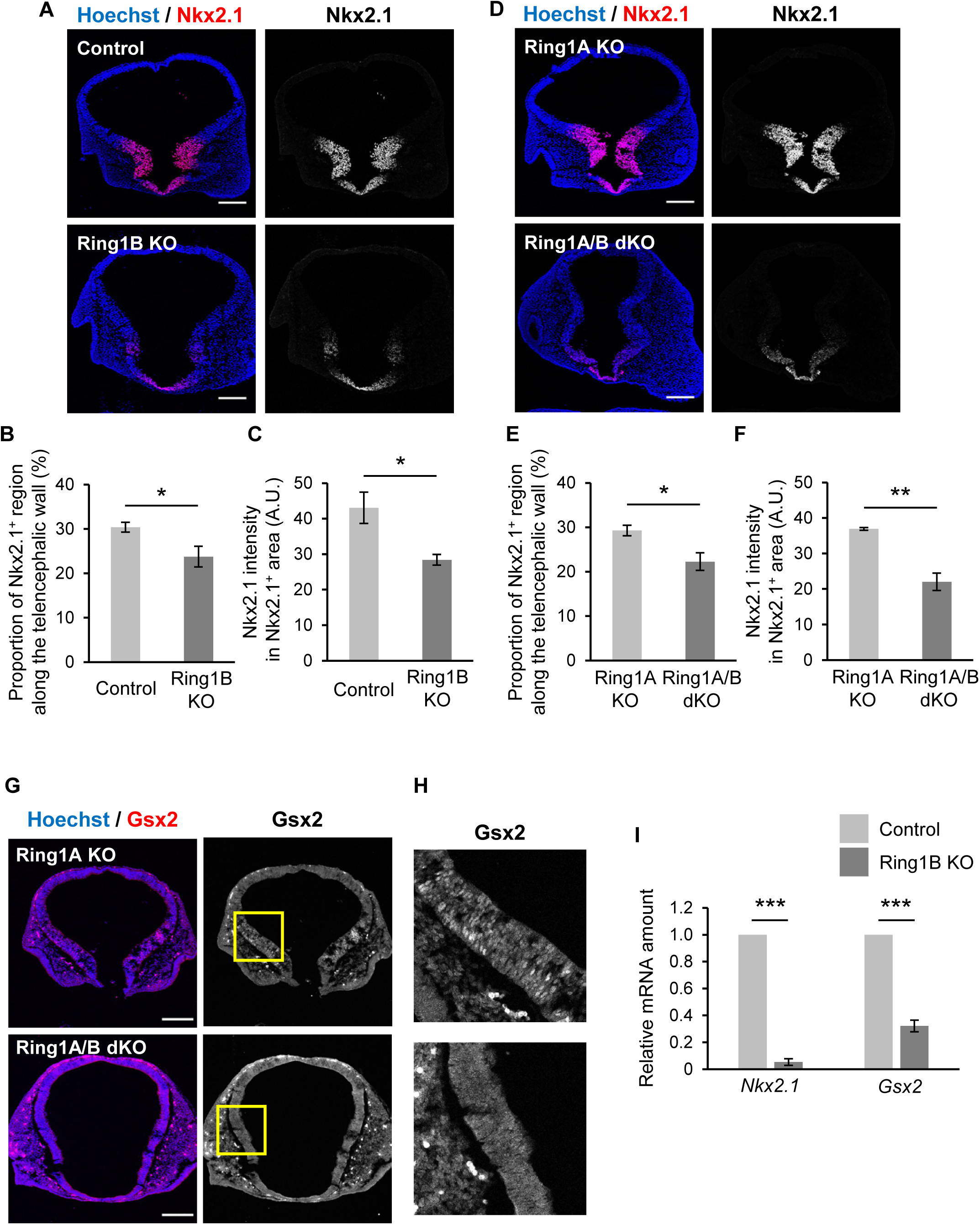
Deletion of *Ring1* down-regulates the expression of ventral-specific transcription factors in NPCs of the ventral telencephalon. (A, D, G, H) Coronal sections of the brain of control or Ring1B KO mice (A) or of Ring1A KO or Ring1A/B dKO mice (D, G) at E10 were subjected to immunohistofluorescence staining with antibodies to Nkx2.1 (A, D) or to Gsx2 (G). Nuclei were counterstained with Hoechst 33342. Boxed regions (300 by 300 µm) in (G) are shown at higher magnification in (H). Scale bars, 200 µm. (B, C, E, F) Quantification of immunostaining intensity for Nkx2.1 in images similar to those in (A) and (D). The Nkx2.1^+^ perimeter length as a proportion of total perimeter length for the telencephalic wall was determined (B, E). The average Nkx2.1 immunostaining intensity within the Nkx2.1^+^ region (pixels) was also determined in arbitrary units (A.U.) for each section, with multiple sections of the telencephalon along the rostrocaudal axis being examined for each embryo (C, F). Data are means ± s.e.m. of values for three embryos. *P < 0.05, **P < 0.01 (two-tailed Student’s unpaired *t* test). (I) Reverse transcription (RT) and quantitative polymerase chain reaction (qPCR) analysis of relative *Nkx2.1* or *Gsx2* mRNA abundance normalized to β-actin mRNA in CD133^+^ NPCs isolated from the GE of control or Ring1B KO mice at E11. Data are means ± s.e.m. of averaged values for four litters (4-7 embryos per litter). *** P < 0.001 (two-tailed Student’s paired *t* test).

We also examined the expression of Gsx2, which is highly enriched in the LGE and whose mRNA is present in both the LGE and MGE (Toresson et al., 2000). Immunostaining indeed revealed the expression of Gsx2 protein within nuclei of NPCs in the LGE of control mice at E10, whereas such expression was markedly attenuated in Ring1A/B dKO mice (Figure 2G, H). Moreover, *Ring1b* deletion significantly reduced the level of *Gsx2* mRNA in CD133^+^ NPCs isolated from the GE of the telencephalon at E11 (Figure 2I). These results together indicated that Ring1 plays a role in expression of ventral transcription factors in the ventral region of the telencephalon.

### Deletion of *Ring1* increases the expression of CTX-specific transcription factors in NPCs of the ventral telencephalon

We then investigated whether deletion of *Ring1* affects the expression of dorsal-specific transcription factors in the developing telencephalon. Pax6 contributes to development of the CTX, with its expression being restricted to the dorsal pallium in mice (Corbin et al., 2003). However, we found that the expression of Pax6 extended to the ventral region of the telencephalon in Ring1B KO mice as well as in Ring1A/B dKO mice (Figure 3A, B, data not shown). Indeed, the dorsoventral gradient of Pax6 expression was shallower in Ring1B KO mice than in control mice (Figure 3C). We also examined the role of Ring1 in regulation of Emx1, another CTX-specific transcription factor (Simeone et al., 1992). Deletion of *Ring1b* increased the amount of *Emx1* mRNA in CD133^+^ NPCs isolated from the GE of the telencephalon at E11 (Figure 3D). These results together indicated that Ring1 suppresses the expression of dorsal transcription factors in the ventral region of the telencephalon and thus prevents “dorsalization” of this region during the early stage of development. Of note, deletion of *Ring1b* with the use of the *Foxg1-IRES-Cre* transgene (that is, from ∼E9.0) did not appear to promote dorsalization of the ventral telencephalon (data not shown), revealing a time window for sensitivity to Ring1-dependent D-V regionalization.

**Figure 3.**
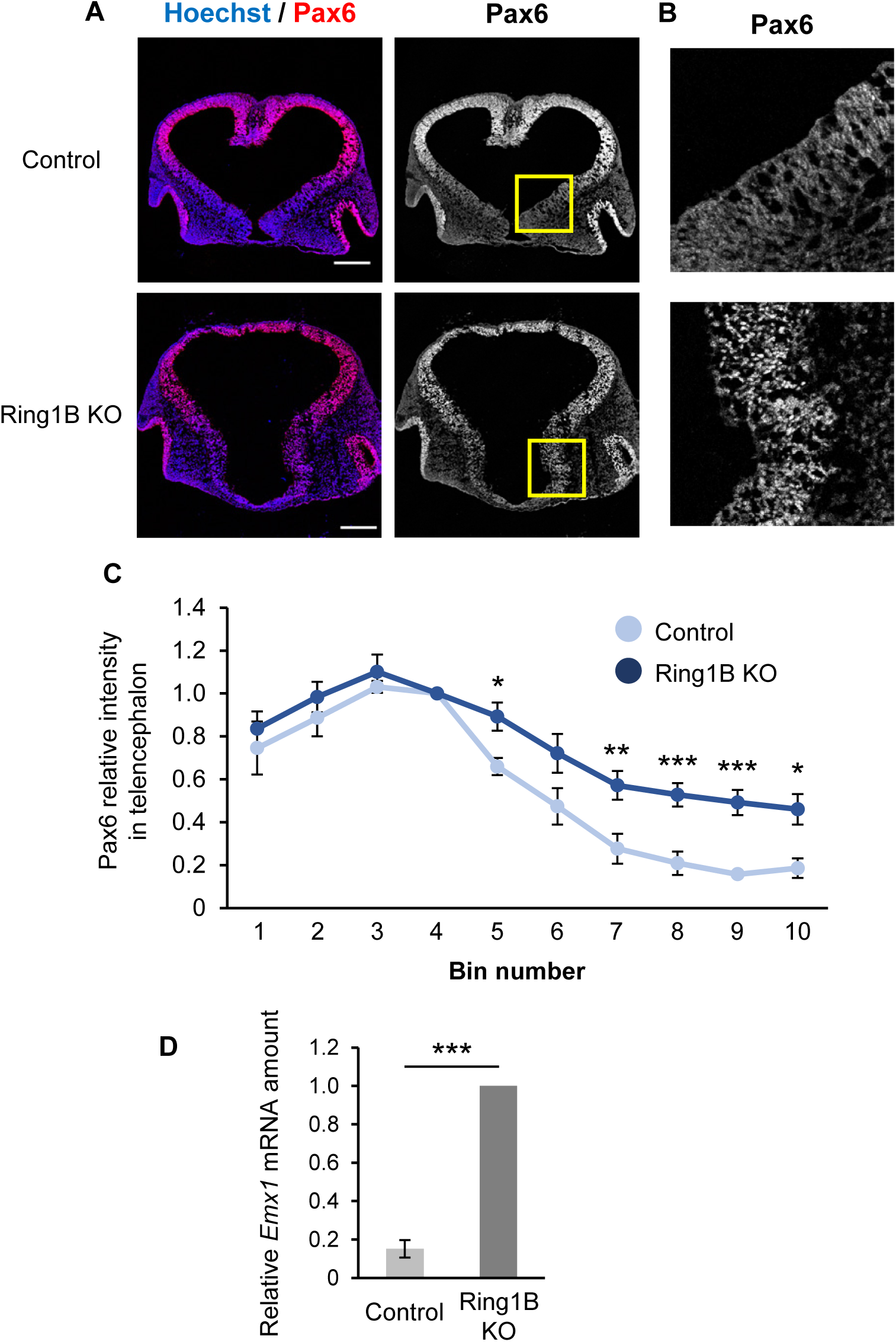
Deletion of *Ring1* increases the expression of CTX-specific transcription factors in NPCs of the ventral telencephalon. (A, B) Coronal sections of the brain of control or Ring1B KO mice at E10 were subjected to immunohistofluorescence staining with antibodies to Pax6. Nuclei were counterstained with Hoechst 33342. Boxed regions (300 by 300 µm) in (A) are shown at higher magnification in (B). Scale bars, 200 µm. (C) Quantification of immunostaining intensity of Pax6 for images similar to those in (A). The telencephalic wall was divided into 10 bins, from 1 (dorsal) to 10 (ventral), and the average immunostaining intensity of Pax6 was determined in each bin and normalized by the average value for bin 4. The average intensity was determined from multiple sections of the telencephalon along the rostrocaudal axis for each embryo. Data are means ± s.e.m. of values from three embryos. *P < 0.05, **P < 0.01, ***P < 0.001 versus the corresponding value for control mice (two-tailed Student’s unpaired *t* test). (D) RT-qPCR analysis of relative *Emx1* mRNA abundance normalized to β-actin mRNA in CD133^+^ NPCs isolated from the ventral telencephalon of control or Ring1B KO mice at E11. Data are means ± s.e.m. of averaged values for four litters (4-7 embryos per litter). *** P < 0.001 (two-tailed Student’s paired *t* test).

### Deletion of *Ring1* confers dorsalized expression patterns of proneural genes in the ventral telencephalon

Given that *Ring1* deletion appeared to induce dorsalization of the expression patterns of transcription factors related to NPC specification along the D-V axis, we examined the expression of proneural genes that contribute to region-specific neuronal differentiation. The basic helix-loop-helix proteins Neurog1 and Ascl1 are pallium- and subpallium-specific proneural factors, respectively (Casarosa et al., 1999; Fode et al., 2000). There was thus little overlap of Neurog1 and Ascl1 expression at the pallium-subpallium boundary of control mice at E10 (Figure 4A). However, deletion of *Ring1b* resulted in a ventral shift of the ventral border of Neurog1 expression and a marked overlap of Neurog1 expression with Ascl1 expression in the ventral region (Figure 4A–C, Figure S2), again suggesting that loss of Ring1 induces dorsalization of the early-stage telencephalon. Deletion of *Ring1b* did not obviously shift the dorsal border of Ascl1 expression (Figure 4C) but significantly reduced the level of Ascl1 within the LGE (Figure 4D). In Ring1A/B dKO mice, the border of the Neurog1^+^ region was also shifted ventrally (Figure 4E), and the level of Ascl1 protein in the ventral region appeared to be reduced to a greater extent than in Ring1A KO or Ring1B KO mice (Figure 4F). These results supported the notion that Ring1 plays a pivotal role in the establishment of ventral identity in the early stage of telencephalic development.

**Figure 4.**
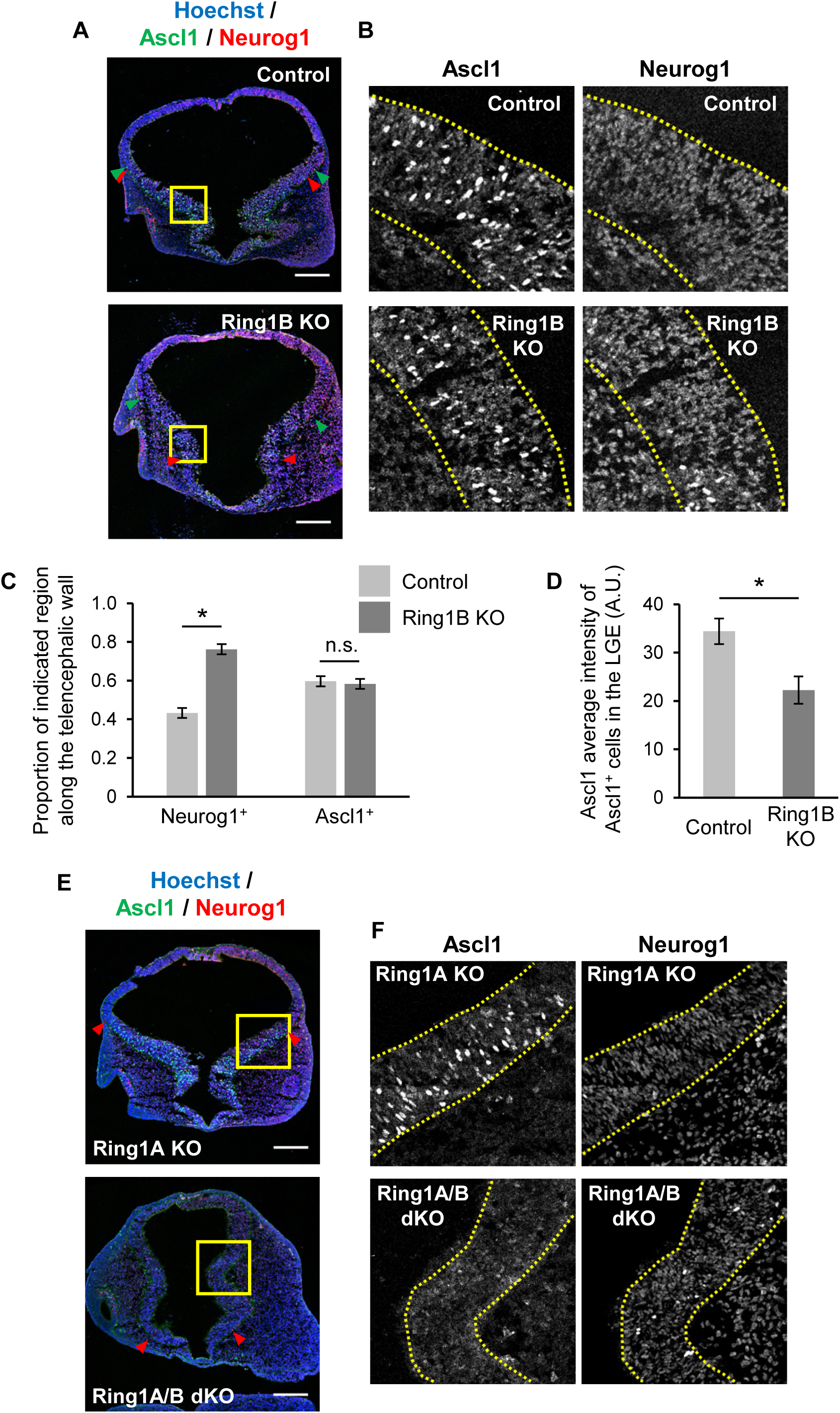
Deletion of *Ring1* confers dorsalized expression patterns of proneural genes in the ventral telencephalon. (A, B, E, F) Coronal sections of the brain of control or Ring1B KO mice (A) or of Ring1A KO or Ring1A/B dKO mice (E) at E10 were subjected to immunohistofluorescence staining with antibodies to Neurog1 and to Ascl1. Nuclei were counterstained with Hoechst 33342. The boxed regions (200 by 200 µm or 300 by 300 µm) in (A) and (E) are shown at higher magnification in (B) and (F), respectively. Green and red arrowheads represent the dorsal and ventral borders of Ascl1^+^ and Neurog1^+^ regions, respectively. The telencephalic wall is demarcated by yellow dotted lines. Note that the Ascl1^+^Neurog1^+^ region was enlarged by *Ring1b* deletion. Scale bars, 200 µm. (C, D) Neurog1^+^ perimeter length or Ascl1^+^ perimeter length was determined as a proportion of total perimeter length for the telencephalic wall of control and Ring1B KO mice (C). The average of Ascl1 immunostaining intensity within Ascl1^+^ cells in the LGE (pixels) was also determined for each section and then corrected for the intensity in the dorsal telencephalon (D). The average intensity of multiple sections of the telencephalon along the rostrocaudal axis was determined for each embryo. Data are means ± s.e.m. of values from three embryos. *P < 0.05 (two-tailed Student’s paired *t* test).

### Ring1 suppresses BMP and Wnt signaling pathways in the early-stage ventral telencephalon

We next investigated whether *Ring1* deletion affects the gene expression profile of NPCs in the ventral telencephalon. Transcripts isolated from CD133^+^ NPCs derived from the GE of the control or Ring1B KO telencephalon at E11 were subjected to RT, and the resulting cDNA was amplified with the use of the Quartz protocol (Sasagawa et al., 2013) and subjected to high-throughput sequencing analysis (Figure 5A). Differentially expressed genes were determined with the use of *edgeR* of the *R* package (McCarthy et al., 2012; Robinson et al., 2010). We identified more up-regulated genes (953) than down-regulated genes (238) in ventral NPCs from Ring1B KO mice compared with those from control mice (Figure 5B, C), consistent with the general role of PcG proteins in gene repression. Importantly, the expression of genes for dorsal-specific transcription factors such as *Emx1*, *Emx2*, and *Msx1* was up-regulated, whereas that of genes for ventral-specific transcription factors such as *Nkx2.1* and *Olig2* (Lu et al., 2000; Takebayashi et al., 2000) was down-regulated, in the NPCs from Ring1B KO mice (Figure 5B, C, Supplementary Table 1). These results thus confirmed the role of Ring1 in suppression of the dorsalization of ventral NPCs.

**Figure 5.**
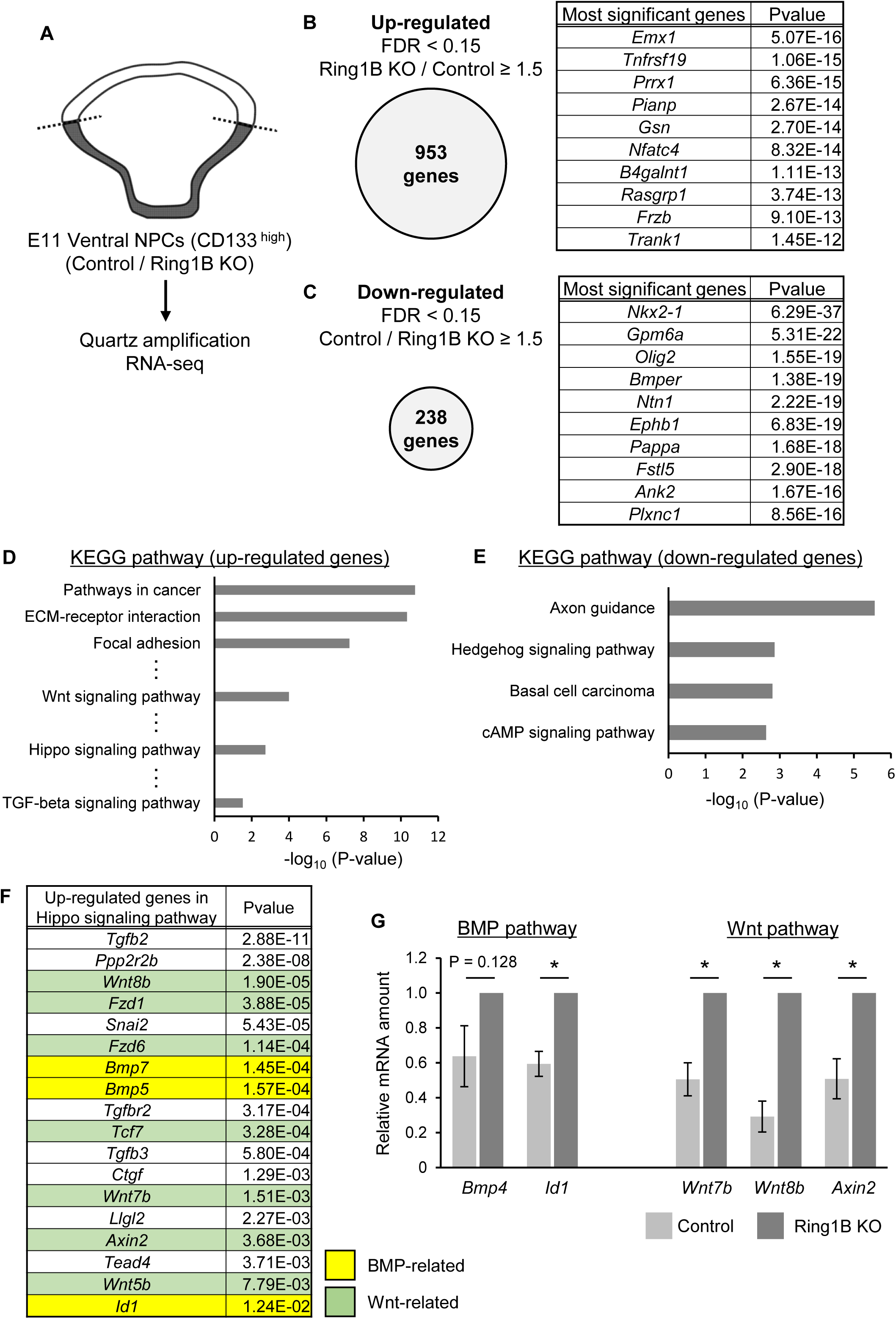
Genome-wide gene expression analysis of Ring1B-null NPCs derived from the ventral telencephalon. (A) Total RNA isolated from CD133^+^ NPCs derived from the GE of control or Ring1B KO mice at E11 was subjected to RT, and the resulting cDNA was amplified by the Quartz method and subjected to high-throughput sequencing analysis. A total of three samples prepared from 1, 1, or 2 embryos was analyzed for each genotype. (B, C) Genes whose expression was up-regulated (B) or down-regulated (C) in NPCs of Ring1B KO mice were defined as those whose Ring1B KO/control or control/Ring1B KO fold change, respectively, was ≥ 1.5, with a false discovery rate (FDR) of < 0.15 (left). The list of genes with the 10 lowest P values in each category is also shown (right). (D, E) Enriched pathways among up-regulated genes (D) and down-regulated genes (E) were determined by KEGG pathway analysis. For the full list of differentially expressed genes and enriched pathways, see Supplementary Table 1. (F) Up-regulated genes categorized to the Hippo signaling pathway include genes related to the BMP signaling pathway (highlighted in yellow) or the Wnt signaling pathway (highlighted in green). (G) RT-qPCR analysis of the relative abundance of *Bmp4*, *Id1*, *Wnt7b*, *Wnt8b*, and *Axin2* mRNAs normalized to β-actin mRNA in NPCs of Ring1B KO or control mice. Data are means ± s.e.m. for three or four independent experiments. *P < 0.05 (two-tailed Student’s paired *t* test).

To shed light on the mechanism by which Ring1 establishes (or maintains) ventral identity in ventral telencephalic NPCs, we performed KEGG (Kyoto Encyclopedia of Genes and Genomes) pathway analysis for both the up-regulated and down-regulated gene sets (Figure 5D, E). Among the genes whose expression was up-regulated by *Ring1b* deletion, pathway analysis revealed an enrichment of categories such as pathways in cancer, extracellular matrix (ECM)–receptor interaction and focal adhesion (Figure 5D). This enrichment may be due in part to derepression of protocadherin-γ and collagen family genes in Ring1B-deficient NPCs compared with control NPCs (Supplementary Table 1). Furthermore, we found that Hippo, Wnt, and transforming growth factor–β (TGF-β) signaling pathways were also enriched among the genes whose expression was up-regulated by *Ring1b* deletion (Figure 5D). The up-regulated genes categorized in the Hippo signaling pathway included genes related to BMP and Wnt signaling pathways (Figure 5F). Of interest, the BMP inhibitor gene *Bmper* was among the top 10 down-regulated genes (Figure 5C). RT-qPCR analysis also showed that the expression of *Bmp4* and *Id1*, which are major components of BMP signaling, as well as that of *Wnt7b*, *Wnt8b*, and *Axin2*, which are major components of Wnt signaling, were increased in Ring1B-null NPCs compared with control NPCs (Figure 5G). These results together indicated that Ring1B suppresses BMP and Wnt signaling pathways in NPCs of the ventral telencephalon at E11.

We also monitored the activity of the Wnt signaling pathway by examining the distribution of *Axin2* mRNA with *in situ* hybridization analysis. The highest level of *Axin2* expression is normally confined to the dorsal midline (presumptive cortical hem), with a lower level of expression also occurring in the pallium (presumptive neocortex) of the early-stage telencephalon. However, the region showing the greatest abundance of *Axin2* mRNA at E10 was expanded in Ring1A/B dKO mice compared with Ring1A KO mice (Figure 6A, B), suggesting that Ring1 suppresses the activity of Wnt signaling outside of the dorsal midline in the telencephalon, with the exception of the most ventral region. We also examined the distribution of Id1 protein as a marker of the activity of the BMP signaling pathway. The highest level of Id1 expression is normally confined to the dorsal midline in the early-stage telencephalon, and again deletion of *Ring1a* and *Ring1b* resulted in expansion of this region to the entire ventricular wall of the telencephalon at E10 (Figure 6C–E). These results suggested that Ring1 suppresses the expression of Id1, and therefore possibly the activity of BMP signaling, outside of the dorsal midline at the early stage of telencephalic development.

**Figure 6.**
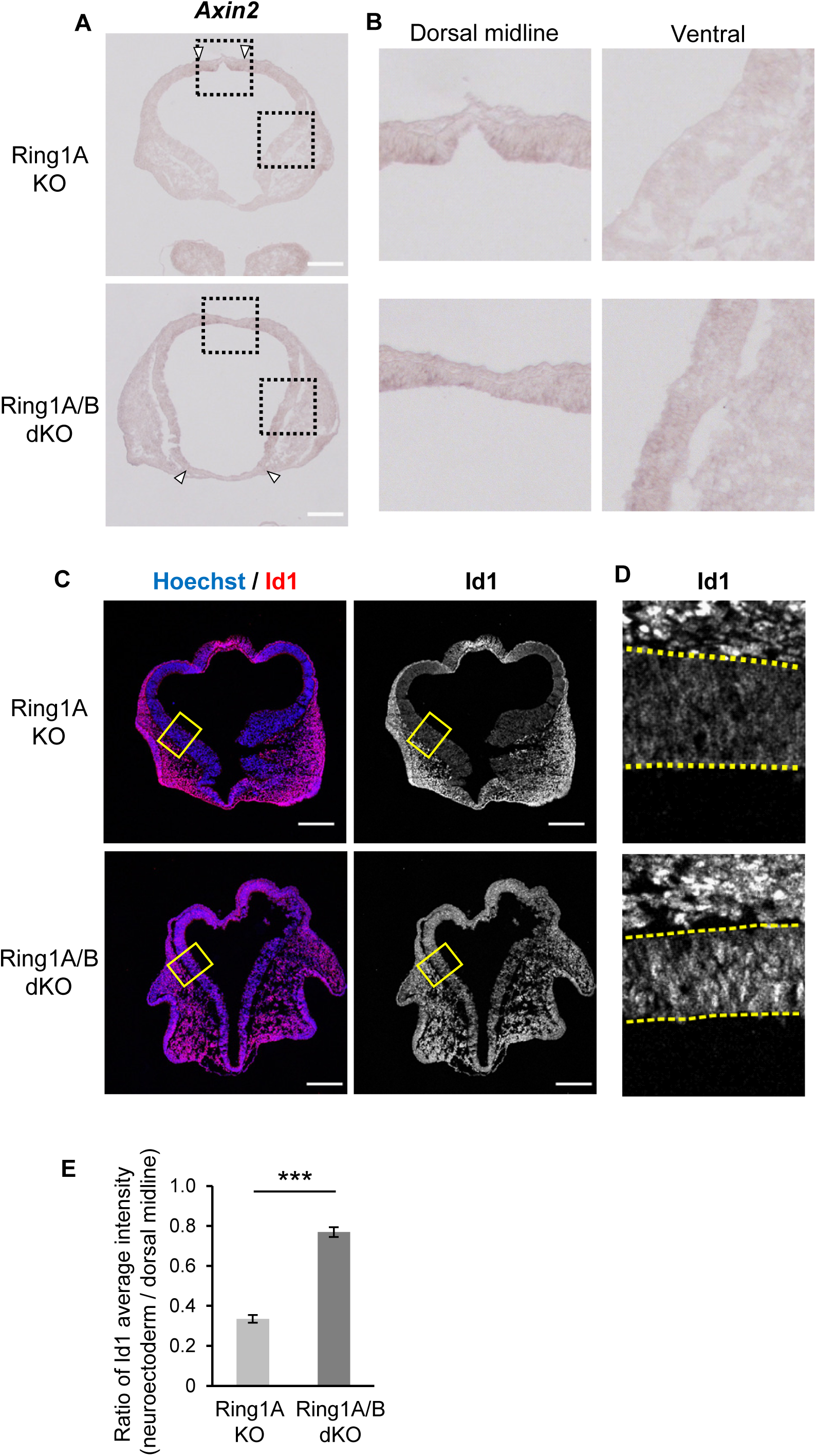
Deletion of *Ring1* activates BMP and Wnt signaling pathways in the early-stage telencephalon. (A, B) Coronal sections of the brain of Ring1A KO or Ring1A/B dKO mice at E10 were subjected to *in situ* hybridization analysis of *Axin2* mRNA. Boxed regions (300 by 300 µm) in (A) are shown at higher magnification in (B). Open arrowheads represent the boundaries of Axin2^high^ regions. Scale bars, 200 µm. (C, D) Coronal sections of the brain of Ring1A KO or Ring1A/B dKO mice at E10 were subjected to immunohistofluorescence staining with antibodies to Id1. Boxed regions (150 by 200 µm) in (C) are shown at higher magnification in (D). Nuclei were counterstained with Hoechst 33342. The neural tissue (telencephalic wall) is outlined by yellow dotted lines. Scale bars, 200 µm. (E) Quantification of Id1 immunostaining intensity in the telencephalon inside and outside of the dorsal midline. The average intensity of Id1 signals in the dorsal 10% and ventral 90% of the telencephalic wall was determined from images similar to those in (C). The average of multiple sections of the telencephalon along the rostrocaudal axis was determined as a representative score for each embryo. Data are means ± s.e.m. of values for three embryos of each genotype. ***P < 0.001 (two-tailed Student’s unpaired *t* test).

### Ring1 promotes *Shh* expression and activates the Shh signaling pathway in the early-stage telencephalon

In contrast to the genes whose expression was up-regulated in Ring1B-null NPCs, KEGG pathway analysis revealed an enrichment of hedgehog signaling pathway among the down-regulated genes (Figure 5E). Consistent with this observation, we found that the expression levels of genes related to Shh signaling—such as *Gli1*, *Gli2*, *Ptch1*, and *Ptch2*—were significantly lower in ventral NPCs from Ring1B KO mice compared with those from control mice at E11 (Figure 7A, B), suggesting that Ring1B is essential for activation of Shh signaling in ventral NPCs at this early stage of telencephalic development.

**Figure 7.**
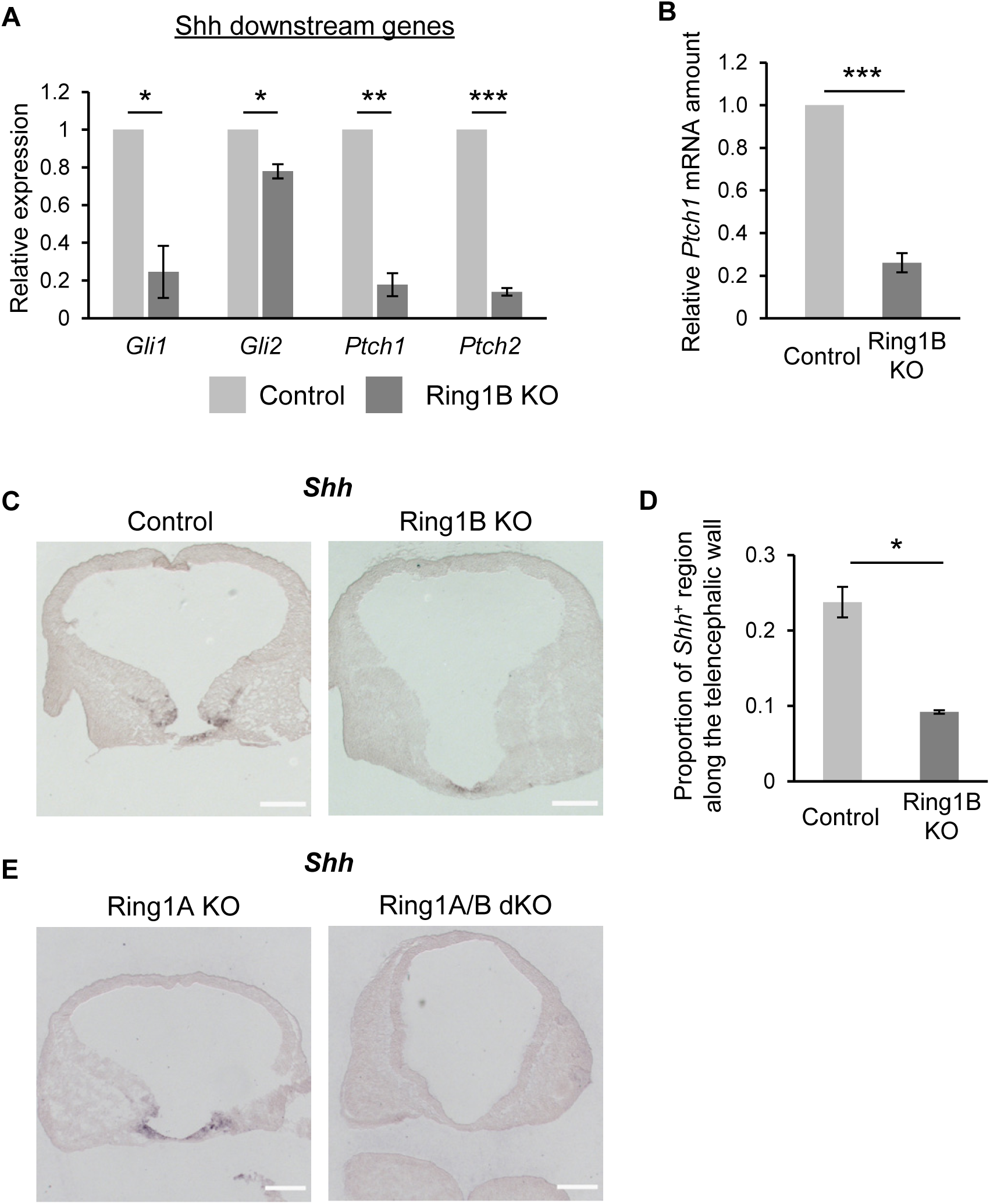
Deletion of *Ring1* attenuates the expression of *Shh* in the telencephalon. (A) The RPKM (reads per kilobase of mRNA model per million total reads) scores for *Gli1*, *Gli2*, *Ptch1*, and *Ptch2* in the RNA-sequencing analysis shown in Figure 5A were normalized by those for the corresponding control sample in each experiment. Data are means ± s.e.m. of values from three experiments. *P < 0.05, **P < 0.01, ***P < 0.001 (two-tailed Student’s paired *t* test). (B) RT-qPCR analysis of relative *Ptch1* mRNA abundance normalized to β-actin mRNA in ventral NPCs of Ring1B KO or control mice at E11. Data are means ± s.e.m. of values from four independent experiments. ***P < 0.001 (two-tailed Student’s paired *t* test). (C, E) Coronal sections of the brain of control or Ring1B KO mice (C) or of Ring1A KO or Ring1A/B dKO mice (E) at E10 were subjected to *in situ* hybridization analysis of *Shh* mRNA. Scale bars, 200 µm. (D) The *Shh*^+^ perimeter length as a proportion of total perimeter length for the telencephalic wall determined from sections similar to those in (C). The average *Shh*^+^ signal intensity for multiple sections of the telencephalon along the rostrocaudal axis was determined for each embryo. Data are means ± s.e.m. of values for three embryos of each genotype. *P < 0.05 (two-tailed Student’s unpaired *t* test).

Given the attenuated expression of Shh target genes in Ring1B-deficient ventral NPCs, we examined whether deletion of *Ring1b* might affect the expression of *Shh*. *In situ* hybridization analysis of mice at E10 revealed that *Ring1b* deletion markedly reduced the abundance of *Shh* mRNA (Figure 7C, D), which is normally found at the ventral midline (presumptive preoptic area) of the developing telencephalon at this stage (Figure 7C). Furthermore, *Shh* expression was not apparent in the telencephalon of Ring1A/B dKO embryos (Figure 7E). These results together indicated that Ring1 is required for expression of Shh, the major ventral morphogen, which might explain the overall dorsalization phenotype of the Ring1-deficient telencephalon.

It remained unclear, however, whether the down-regulation of Shh target gene expression induced by *Ring1* deletion was due simply to the attenuation of *Shh* expression or was also due to an inability of NPCs to express these genes in response to Shh signaling. We therefore prepared *in vitro* cultures of telencephalic NPCs at E10 and examined their responsiveness to Shh signaling by the addition of a Smoothened agonist (SAG) for 24 h. The extent of the induction of the Shh target genes *Gli1* and *Ptch1* was similar for NPCs isolated from Ring1B KO mice and from control mice (Figure 8A–C), suggesting that *Ring1b* deletion did not substantially affect the regulation of these Shh target genes and that the regulation of *Shh* expression itself is crucial for Ring1-dependent ventral identity.

**Figure 8.**
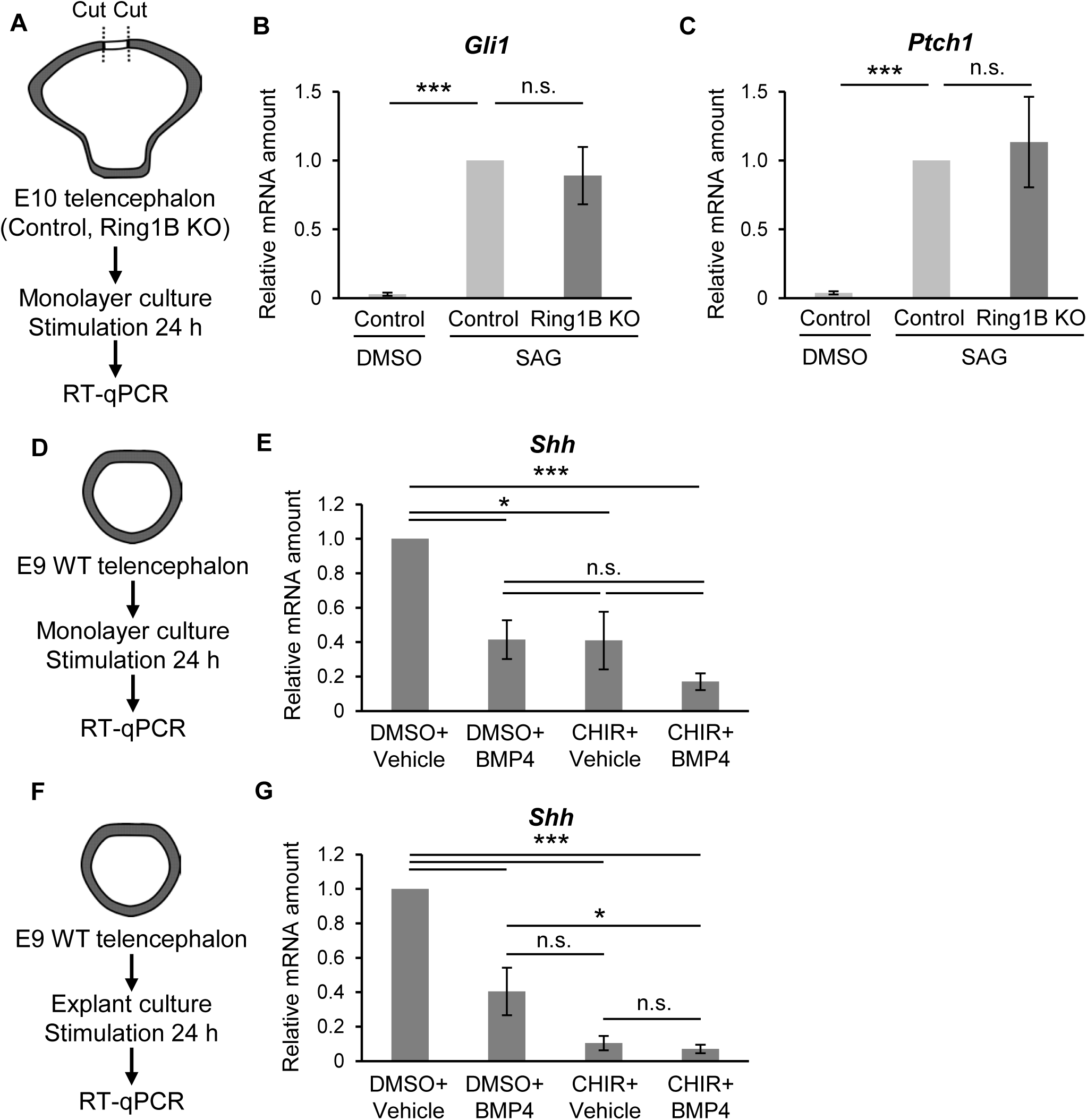
**Forced activation of the BMP and Wnt pathways inhibits *Shh* expression in early-stage telencephalic NPCs** (A) NPCs isolated from the telencephalon (outside of the dorsal midline) of control or Ring1B KO mice at E10 were cultured as monolayers for 6 h, exposed for an additional 24 h to 2 µM Smoothened agonist (SAG) or dimethyl sulfoxide vehicle, and then subjected to RT-qPCR analysis. (B, C) RT-qPCR analysis of relative *Gli1* (B) and *Ptch1* (C) mRNA abundance normalized to β-actin mRNA in NPCs treated as in (A). Data are means ± s.e.m. for three independent experiments. ***P < 0.001 (one-way ANOVA followed by Bonferroni’s multiple-comparison test). (D, F) The telencephalon of WT (ICR) mice at E9 was subjected to dissociation for monolayer culture (D) or was cultured as an explant (F) for 6 h before exposure for an additional 24 h to BMP4 (50 ng/ml) or 5 µM CHIR-99021 as indicated followed by RT-qPCR analysis. (E, G) RT-qPCR analysis of relative *Shh* mRNA abundance normalized to β-actin mRNA in cultures as in (D) and (E), respectively. Data are means ± s.e.m. for four independent experiments. *P < 0.05, ***P < 0.001 (one-way ANOVA followed by Bonferroni’s multiple-comparison test).

How does Ring1 promote *Shh* expression? Given the general role of PcG proteins in gene repression, it was plausible that Ring1 indirectly increases *Shh* expression through repression of genes whose products inhibit *Shh* expression. Implantation of beads soaked with recombinant BMP in the anterior neuropore of chick embryos was previously shown to inhibit *Shh* and *Nkx2.1* expression (J. A. Golden et al., 1999; Ohkubo et al., 2002). Furthermore, forced activation of canonical Wnt signaling by expression of a stabilized form of β-catenin was found to result in repression of ventral marker genes such as *Nkx2.1* in the mouse subpallium (Backman et al., 2005). It was therefore possible that activation of BMP and Wnt signaling pathways might account for the down-regulation of *Shh* expression in Ring1-deficient mice, although it remained unclear whether Wnt signaling alone is able to regulate the level of *Shh* expression. We therefore examined whether activation of Wnt signaling can reduce the level of *Shh* mRNA in collaboration with BMP signaling in dissociated (monolayer) cultures and explant cultures prepared from the telencephalon of WT mice at E9. The addition of an activator of canonical Wnt signaling (the glycogen synthase kinase 3 inhibitor CHIR-99021) indeed significantly reduced the abundance of *Shh* mRNA in the dissociated telencephalic culture and, to a greater extent, in the explant culture (Figure 8D–G) under the condition. Moreover, exposure to both BMP4 and CHIR-99021 tended to have a greater effect on the amount of *Shh* mRNA in the dissociated culture than did either agent alone (Figure 8E, G). The activation of BMP and Wnt signaling pathways may therefore cooperate to suppress the expression of *Shh* in the telencephalon at this early stage of development, consistent with the notion that the Ring1-dependent establishment of ventral identity is mediated by suppression of these signaling pathways.

### BMP and Wnt ligand genes are direct targets of PcG proteins in early-stage telencephalic NPCs

Given the dysregulation of BMP and Wnt ligand gene expression induced by *Ring1* deletion, we next examined whether these genes are direct targets of PcG proteins by performing chromatin immunoprecipitation (ChIP)–qPCR assays for H3K27me3, a histone modification catalyzed by PRC2, as well as for Ring1B with the telencephalon of WT mice at E9. We indeed detected significant or nearly significant deposition of H3K27me3 at the promoters of *Bmp4*, *Bmp7*, *Wnt7b*, and *Wnt8b* at levels similar to those apparent at the promoters of *Hoxa1* and *Hoxd3*, which were examined as positive controls (Figure 9B). Ring1B was found to be enriched at the promoters of *Bmp4* and *Wnt8b*, but not at those of *Bmp7* and *Wnt7b* (Figure 9C), suggesting that PcG proteins directly regulate the expression of at least *Bmp4* and *Wnt8b* in the early-stage telencephalon.

**Figure 9.**
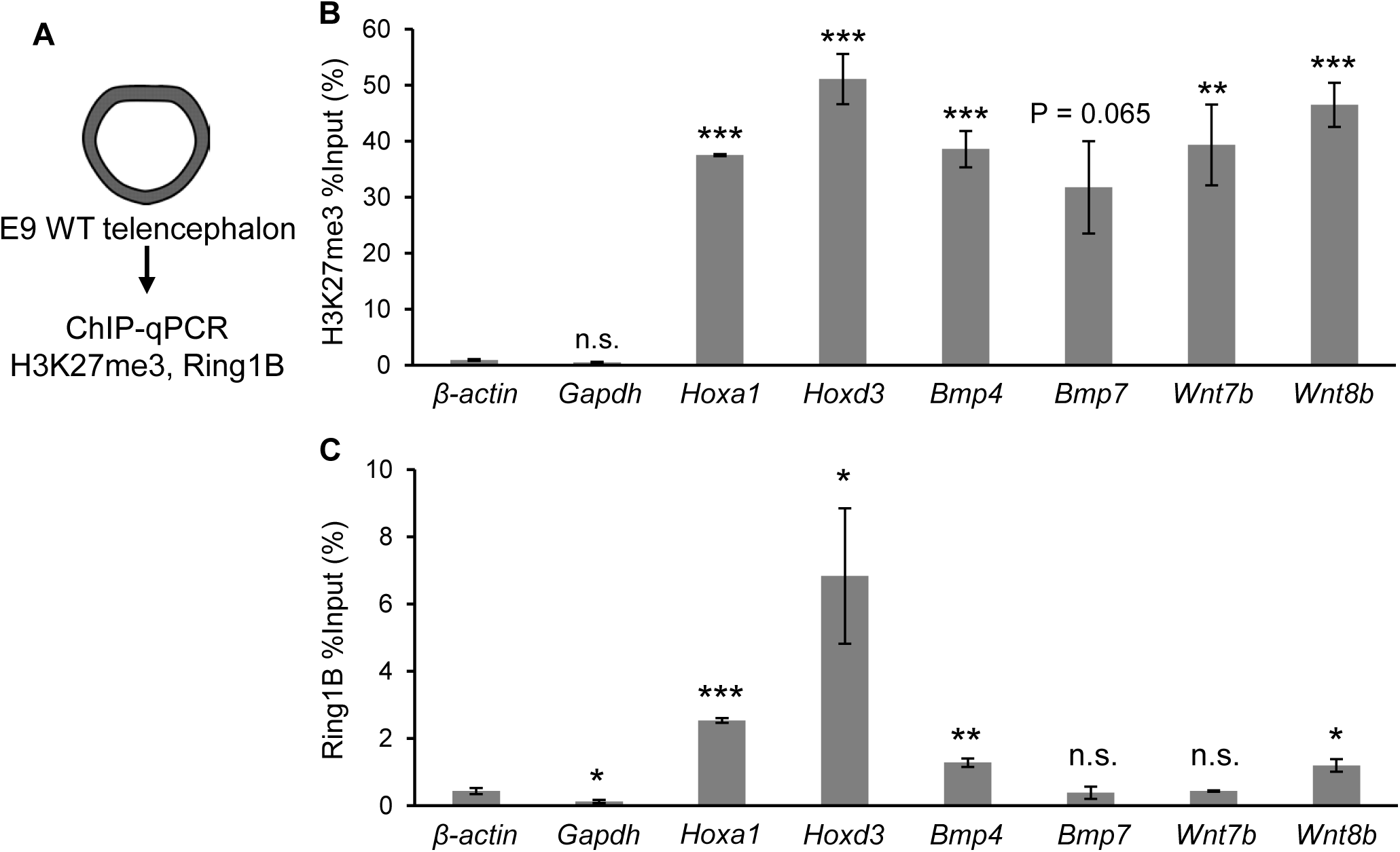
H3K27me3 deposition and Ring1B binding to the promoters of *Bmp4* and *Wnt8b* in early-stage telencephalic NPCs. (A) The telencephalon was isolated from WT (ICR) mice at E9 for ChIP-qPCR analysis with antibodies to H3K27me3 or to Ring1B. (B, C) ChIP-qPCR analysis of H3K27me3 deposition (B) and Ring1B binding (C) at the indicated promoters. *Hoxa1* and *Hoxd3* were examined as positive controls, and *Gapdh* and the β-actin gene as negative controls. Data are expressed as the percentage to input value and are means ± s.e.m. for three independent experiments. *P < 0.05, **P < 0.01, *** P < 0.001 versus the corresponding value for the β-actin gene (two-tailed Student’s unpaired *t* test).

## Discussion

Morphogenetic signals and their downstream transcription factors determine regional identity along the D-V axis in the developing telencephalon. Mutual inhibition between such signaling plays a pivotal role in segregation of regional identity, but the contribution of epigenetic mechanisms that control the permissiveness for transcriptional activation to this telencephalic D-V regionalization have remained largely unknown. We have now found that Ring1A and Ring1B, core components of PRC1, play an essential role in establishment of the spatial expression patterns of morphogenetic signals and transcription factors along the D-V axis and in consequent regionalization of the telencephalon at the early stage of development. Our results thus indicate that Ring1 is required for expression of *Shh* in the ventral telencephalon and that the ablation of *Ring1* results in dorsalization of the ventral telencephalon. This dorsalization phenotype of the Ring1-deficient telencephalon is likely due in part to down-regulation of the ventralizing morphogen Shh, given that the inactivation of *Shh* gives rise to morphological and molecular phenotypes similar to those associated with *Ring1* deletion, including the induction of a rounder shape and malformation of the dorsal midline in the telencephalic wall (Chiang et al., 1996; Rallu et al., 2002), increased apoptosis (Aoto et al., 2009), and attenuated expression of ventral transcription factors such as Nkx2.1, Gsx2, and Ascl1 (Blaess et al., 2014). Moreover, our results indicate that Ring1 is required for suppression of BMP and Wnt signaling pathways and that genes for BMP and Wnt ligands are direct targets of Ring1 and other PcG proteins. Together with the observations that ectopic activation of BMP signaling (J. A. Golden et al., 1999; Ohkubo et al., 2002) or Wnt signaling (this study) is able to suppress the transcription of *Shh* in the developing telencephalon, these results suggest that Ring1 suppresses BMP and Wnt signaling in the telencephalon (outside of the dorsal midline) and thereby generates a permissive state for *Shh* expression, which is essential for establishment of ventral identity (Figure 10). In addition to the activation of *Shh* expression, suppression of BMP and Wnt signaling pathways *per se* may contribute to the Ring1-mediated establishment of ventral identity by a Shh-independent mechanism (Figure 10).

**Figure 10.**
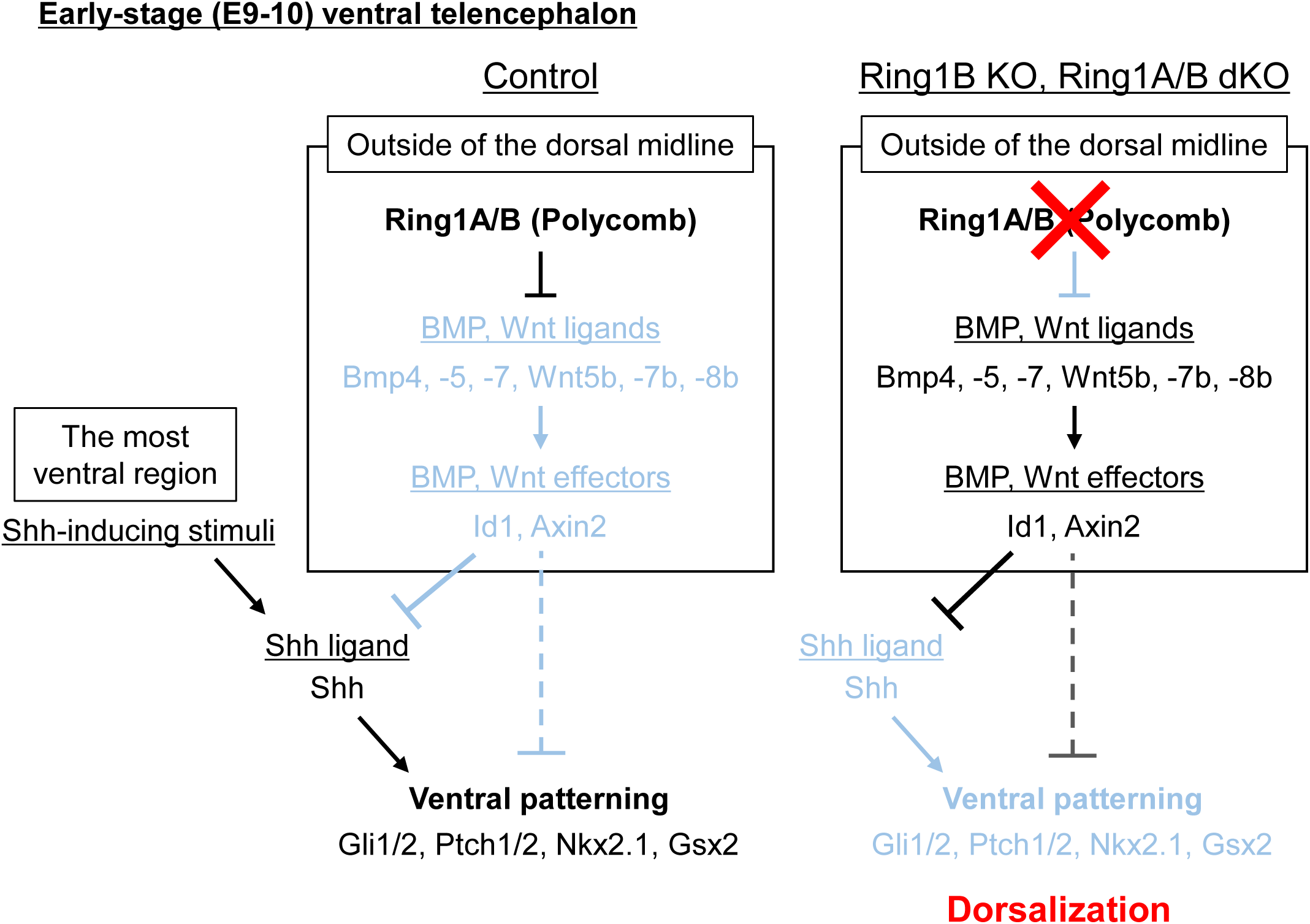
Schematic summary of Ring1-mediated ventral specification in the early-stage mouse telencephalon. Ring1 suppresses BMP and Wnt signaling pathways outside of the dorsal midline at the early stage of mouse telencephalic development and thereby confers a permissive state for *Shh* expression. *Shh*-inducing signals to the most ventral portion of telencephalon thus can induce *Shh* in this region, resulting in the ventral patterning of telencephalon. By contrast, Ring1 ablation derepresses and ectopically activates BMP and Wnt signaling pathways outside of the dorsal midline, and thus suppresses Shh-mediated ventral patterning of telencephalon.

Deletion of *Ring1* not only induced dorsalization of NPC identity in the telencephalon but also resulted in aberrant expression patterns of proneural genes. Ascl1 and Neurog1 are expressed in a mutually exclusive manner in WT embryos, in part as a result of the repression of *Ascl1* expression by Neurog1 and Neurog2 (Fode et al., 2000). However, we found that *Ring1b* deletion resulted in a marked increase in the number of cells positive for both Neurog1 and Ascl1. Given that H3K27me3 and H2AK119ub were previously shown to be deposited at the promoters of *Neurog1* and *Ascl1* in early-stage NPCs (Hirabayashi et al., 2009; Tsuboi et al., 2018; data not shown), PcG proteins may participate in the mutually exclusive inhibition of *Neurog1* and *Ascl1* expression and thereby regulate the segregation of neurogenic properties between NPCs.

Deletion of *Ring1b* with the use of the *Nestin-CreERT2* transgene, which confers Cre expression in the entire CNS at E13.5 (Hirabayashi et al., 2009), or deletion of the gene for the histone methyltransferase Ezh2 with the *Emx1-Cre* transgene, which is expressed in the dorsal telencephalon from E10.5 (Gorski et al., 2002; Pereira et al., 2010), has been shown to induce neurogenesis through derepression of neurogenic genes (Tsuboi et al., 2018). With the use of the *Sox1-Cre* transgene, which is expressed in the neuroepithelium from before E8.5 (Takashima et al., 2007), we have now examined the role of Ring1 in the early stage of telencephalic development, before the onset of the neurogenic phase. During this early stage (for example, at E9), we did not detect promotion of neurogenesis (data not shown), suggesting that a Ring1-indpenedent mechanism is responsible for the suppression of neurogenesis at this time. Of interest, deletion of *Ring1b* with the use of the *Foxg1-IRES-Cre* transgene, which is expressed in the entire telencephalon from ∼E9.0 (Kawaguchi et al., 2016), did not appear to induce dorsalization of the ventral telencephalon (data not shown), suggesting that Ring1-mediated D-V patterning of the telencephalon takes place only during the early stage of development, although the mechanism underlying this temporal restriction remains unclear.

A key related question is how PcG proteins are recruited to specific genes in specific regions of the telencephalon at specific times. Deletion of *Ezh2* in the dorsal midbrain with the use of the *Wnt1-Cre* transgene was previously shown to result in inhibition of Wnt signaling and to promote telencephalic identity at ∼E11.5 (that is, rostralization) (Zemke et al., 2015), in contrast to our finding that *Ring1* deletion activates Wnt signaling in the early-stage telencephalon. This previous study also showed that *Ezh2* deletion increased the expression of *Wif1* and *Dkk2*, both of which encode inhibitors of the Wnt signaling pathway, and that H3K27me3 was deposited at these gene loci in WT embryos, suggesting that PcG proteins contribute to Wnt activation by repressing these Wnt inhibitor genes in the dorsal midbrain. Mechanisms by which recruitment of PcG proteins is regulated in a tissue-, cell type–, or stage-specific manner warrant clarification in future studies.

BMP signaling has been shown to be important for establishment of the dorsal midline and its activity to be confined to this region (Hebert et al., 2002; Panchision et al., 2001; Roy et al., 2014). The relevance of the absence of BMP signaling outside of the dorsal midline has not been known, however. Our results now suggest that suppression of BMP signaling outside of the dorsal midline is required for the expression of *Shh* at the ventral midline and that Ring1 mediates this suppression and thereby sets up a permissive state for *Shh* expression. Given that the suppression of BMP signaling is necessary for neural induction of ectoderm (Wilson and Hemmati-Brivanlou, 1995), its onset may occur before formation of the prospective forebrain. BMP signaling–related targets of PcG proteins identified in embryonic stem cells may be involved in this early process (Shan et al., 2017). The observed increase in Id1 expression in the telencephalon (outside of the dorsal midline) in response to *Ring1* deletion from before E10 suggests that BMP signaling remains repressed in this region but becomes derepressed at the dorsal midline, although the mechanisms underlying this difference remain unknown.

We found that the targets of PcG proteins in the early-stage telencephalon include the genes for BMP and Wnt ligands. Of interest, we detected Ring1B binding to BMP and Wnt ligand gene loci in the telencephalon at E9, but the extent of this binding appeared less than that evident at *Hox* gene loci. In contrast, the levels of H3K27me3 deposition were similar for these two sets of loci. This difference may be due to the operation of different modes of PcG-mediated repression (Tsuboi et al., 2018). Future studies are required to reveal which PcG complexes are responsible for the repression of BMP and Wnt ligand genes.

The robust maintenance of the A-P axis through suppression of *Hox* gene expression by PcG proteins has been well established from flies to mammals (Montavon and Soshnikova, 2014). We now propose that PcG proteins also play an essential role in formation of the D-V axis in the early stage of mouse telencephalic development. Our study thus sheds light on the role of chromatin-level regulation in regionalization of the brain that is dependent on developmental genes that are not necessarily clustered like *Hox* genes.

## Acknowledgments

We thank S. Nishikawa (RIKEN) for providing *Sox1-Cre* mice; I. Nikaido (RIKEN) for providing tips for the Quartz method; K. Imamura, T. Horiuchi, S. Sugano and Y. Suzuki (The University of Tokyo) for performing high-throughput-sequencing analysis; M. Endoh (National University of Singapore) for providing tips for the Ring1B ChIP assay; Y. Koseki (RIKEN) for providing *Ring1* mutant mice; Y. Maeda, R. Nagayoshi, and Y. Kakeya for technical assistance; and members of the Gotoh laboratory for discussion. This research was supported by AMED-CREST of the Japan Science and Technology Agency (grant JP18gm0610013) and by the Ministry of Education, Culture, Sports, Science, and Technology of Japan (JSPS KAKENHI grants JP15H05773, JP16H06481, JP16H06479, and JP16H06279 to Y.G., and JP18K14622, JP19H05253, and JP16H06279 to Y.K.).

## Author Contributions

H.E., Y.K., and Y.G. designed the study and wrote the manuscript. H.E. and Y.K. performed the experiments and analyzed the data. H.K. generated *Ring1a*^−/−^;*Ring1b*^flox/flox^ mice. Y.K. and Y.G. supervised the study.

## Materials and Methods

### Animals

*Ring1b*^flox/flox^ or *Ring1a*^−/−^;*Ring1b*^flox/flox^ mice (Calés et al., 2008; Endoh et al., 2008) were crossed with *Sox1-Cre* transgenic mice (Takashima et al., 2007). Jcl:ICR (CLEA Japan) or Slc:ICR (SLC Japan) mice were studied as WT animals. All mice were maintained in a temperature- and relative humidity–controlled (23° ± 3°C and 50 ± 15%, respectively) environment with a normal 12-h-light, 12-h-dark cycle. They were housed two to six per sterile cage (Innocage, Innovive; or Micro BARRIER Systems, Edstrom Japan) with chips (PALSOFT, Oriental Yeast; or PaperClean, SLC Japan), irradiated food (CE-2, CLEA Japan), and filtered water available ad libitum. Mouse embryos were isolated at various ages, with E0.5 being considered the time of vaginal plug appearance. All animals were maintained and studied according to protocols approved by the Animal Care and Use Committee of The University of Tokyo.

### Plasmid constructs

A pBluescript SK(-) vector encoding mouse Shh was kindly provided by D. Kawaguchi (The University of Tokyo). A portion of the *Axin2* cDNA was cloned by PCR from cDNA derived from the mouse telencephalon and was subcloned into pBluescript SK(-). Amplified sequences are presented in Supplementary table 2.

### Antibodies

Antibodies for immunofluorescence and ChIP analyses included mouse antibodies to Ascl1 (Mash1, BD Pharmingen, 556604, 1:500) and H3K27me3 (MBL, MABI0323, 2 μg/sample for ChIP), goat antibodies to Neurog1 (Santa Cruz, sc-19231, 1:200) and rabbit antibodies to H2AK119ub (Cell Signaling Technology, 8240S, 1:1000), Ring1B (Cell Signaling Technology, 5694S, 1:200, 3 μg/sample for ChIP), Cleaved caspase-3 (Cell signaling Technology, 9664S, 1:1000), Nkx2.1 (TTF1, Abcam, ab76013, 1:1000), Gsx2 (Gsh2, Millipore, ABN162, 1:200), Pax6 (Millipore, AB2237, 1:500) and Id1 (Biocheck, BCH-1/37-2, 1:200). Alexa-labeled secondary antibodies and Hoechst 33342 (for nuclear staining) were obtained from Molecular Probes.

### Immunohistofluorescence analysis

Immunohistofluorescence staining was performed as previously described (Morimoto-Suzki et al., 2014), with minor modifications. In brief, embryos were fixed for 3 h with 4% paraformaldehyde in phosphate-buffered saline (PBS), incubated overnight at 4°C with 30% sucrose in PBS, embedded in OCT compound (Sakura Finetek), and sectioned with a cryostat at a thickness of 10 µm. The sections were exposed to 0.1% Triton X-100 and 3% bovine serum albumin in Tris-buffered saline (blocking solution) for 1 h at room temperature before incubation first overnight at 4°C with primary antibodies diluted in blocking solution and then for 1 h at room temperature with fluorophore-labeled secondary antibodies also diluted in blocking solution. They were finally mounted in Mowiol (Calbiochem) for imaging with a laser-scanning confocal microscope (TSC-SP5, Leica) and ImageJ software (NIH).

### Isolation of ventral NPCs by FACS

The ventral telencephalon was dissected and subjected to enzymatic digestion with a papain-based solution (Sumitomo Bakelite). Cell suspensions were stained with allophycocyanin-conjugated antibodies to CD133 (141210, BioLegend) at a dilution of 1:400 and were then subjected to fluorescence-activated cell sorting (FACS) with a FACSAria instrument (Becton Dickinson). CD133^+^ NPCs were isolated as the top 50% of allophycocyanin-positive cells.

### Quartz-seq analysis

Both cDNA synthesis and amplification were performed with total RNA from 2000 cells as described previously (Sasagawa et al., 2013). In brief, total RNA was purified from cells with the use of Ampure XP RNA (Beckman) and subjected to RT with Super Script III (Thermo Scientific), and the resulting cDNA was purified with Ampure XP (Beckman) and treated with ExoI (Takara) for primer digestion. After addition of a poly(A) tail with terminal deoxynucleotidyl transferase (Roche), the cDNA was subjected to second-strand synthesis and the resulting double-stranded cDNA was amplified with the use of MightyAmp DNA polymerase (Takara). The amplified cDNA was prepared for sequencing with the use of a Nextera XT DNA Sample Prep Kit (Illumina) and subjected to deep sequencing analysis on the Illumina HiSeq2500 platform to yield 36-base single-end reads. Approximately 20 million sequences were obtained from each sample. Sequences were mapped to the reference mouse genome (mm9) with ELAND v2 (Illumina). Only uniquely mapped tags with no base mismatches were used for the analysis. Gene expression was quantitated as reads per kilobase of mRNA model per million total reads (RPKM) on the basis of RefSeq gene models (mm9).

### RT-qPCR analysis

Total RNA was isolated from cells with the use of RNAiso plus (Takara), and up to 0.5 µg of the RNA was subjected to RT with the use of ReverTra Ace qPCR RT Master Mix with gDNA remover (Toyobo). The resulting cDNA was subjected to real-time PCR analysis in a LightCycler 480 instrument (Roche) with either KAPA SYBR FAST for LightCycler 480 (Kapa Biosystems) or Thunderbird SYBR qPCR mix (Toyobo). The amount of each target mRNA was normalized by that of β-actin mRNA. Primer sequences are presented in Supplementary table 2.

### *In situ* hybridization analysis

For preparation of digoxigenin-labeled riboprobes, linearized plasmids containing probe sequences were incubated for 3 h at 37°C with DIG RNA Labeling Mix, Transcription Buffer, and RNA polymerase (Roche) as well as RNase inhibitor (Toyobo). The plasmids were then digested with DNaseI (Takara) for 30 min at 37°C, after which the DNase reaction was stopped by the addition of Stop Solution (Promega). Synthesized riboprobes were purified with the use of a ProbeQuant G-50 column (GE Healthcare) and diluted with hybridization buffer (5× Denhardt’s solution, 5× standard saline citrate, 50% formamide, tRNA at 250 µg/ml, salmon testis DNA at 200 µg/ml, heparin at 100 µg/ml, and 0.1% Tween 20). The riboprobes (0.5 µg/ml) were denatured at 85°C for 5 min, placed on ice for 2 min, and then maintained at 65°C before in situ hybridization. Embryos were fixed for 3 h (*Shh*) or overnight (*Axin2*) with 4% paraformaldehyde in PBS and then incubated with 30% sucrose in PBS, embedded, and sectioned as described for immunohistofluorescence analysis. Sections were fixed for 10 min with 4% paraformaldehyde in PBS, washed with 0.1% Tween 20 in PBS, and incubated at room temperature first with 0.1 M triethanolamine for 3 min and then with the same solution containing 0.1% acetic anhydride for 10 min. They were washed again with 0.1% Tween 20 in PBS before incubation at 65°C first for 1 h with hybridization buffer and then overnight with denatured RNA probes within a humidified box with 50% formamide. The sections were washed twice for 30 min at 65°C with 2× standard saline citrate, twice for 30 min at 65°C with the same solution containing 50% formamide, and three times for 5 min at room temperature with 0.1% Tween 20 in MAB buffer (MABT). After exposure for 1 h at room temperature to 10% fetal bovine serum in MABT, the sections were incubated overnight at 4°C with alkaline phosphatase–conjugated antibodies to digoxigenin (Roche) at a dilution of 1:2000 in the same solution, washed twice for 10 min at room temperature with MABT and twice for 10 min at room temperature with a solution containing 100 mM Tris-HCl (pH 9.5), 100 mM NaCl, 50 mM MgCl_2_, and 0.02% Tween 20, and then incubated at room temperature in the same solution containing NBT-BCIP (nitrotetrazolium blue chloride at 350 µg/ml and 5-bromo-4-chloro-3-indolyl phosphate *p*-toluidine salt at 175 µg/ml) (Roche) until the color appeared. They were finally washed with 0.1% Tween 20 in PBS and mounted in Mowiol (Calbiochem). Images were acquired with an Axiovert 200M microscope fitted with an Axiocam or Axiocam 305 camera (Carl Zeiss) and were processed with ImageJ (NIH).

### ChIP-qPCR analysis

ChIP for Ring1B and H3K27me3 was performed as previously described (Tsuboi et al., 2018), with minor modifications. Cells were fixed with 1% formaldehyde and then suspended in radioimmunoprecipitation (RIPA) buffer for sonication (10 mM Tris-HCl at pH 8.0, 1 mM EDTA, 140 mM NaCl, 1% Triton X-100, 0.1% SDS, 0.1% sodium deoxycholate). They were subjected to ultrasonic treatment to shear genomic chromatin into DNA fragments, and the cell lysates were then diluted with RIPA buffer for immunoprecipitation (50 mM Tris-HCl at pH 8.0, 150 mM NaCl, 2 mM EDTA, 1% Nonidet P-40, 0.1% SDS, 0.5% sodium deoxycholate) and incubated for 1 h at 4°C with ProteinA/G Magnetic Beads (Pierce) to clear nonspecific reactivity. They were then incubated overnight at 4°C with ProteinA/G Magnetic Beads that had previously been incubated overnight at 4°C with antibodies to Ring1B or to H3K27me3. The beads were then isolated and washed three times with wash buffer (2 mM EDTA, 150 mM NaCl, 0.1% SDS, 1% Triton X-100, and 20 mM Tris-HCl at pH 8.0) and then once with wash buffer containing 500 mM NaCl. Immune complexes were eluted from the beads with a solution containing 10 mM Tris-HCl (pH 8.0), 5 mM EDTA, 300 mM NaCl, and 0.5% SDS at 65°C for 15 min, and they were then subjected to digestion with proteinase K (Nakarai) at 37°C for more than 6 h, removal of cross links by incubation at 65°C for more than 6 h, and extraction of the remaining DNA with phenol–chloroform–isoamyl alcohol and ethanol. The DNA was washed with 70% ethanol, suspended in water, and subjected to real-time PCR analysis in a LightCycler 480 instrument (Roche) with Thunderbird SYBR qPCR Mix (Toyobo). Primer sequences are presented in Supplementary table 2.

### Primary culture of the telencephalon and treatment with pharmacological agents

For monolayer culture, primary NPCs were isolated from the indicated regions of the telencephalon of ICR or of Ring1B KO or control mouse embryos. The dissected tissue was thus subjected to digestion with a papain-based solution (Sumitomo Bakelite), and the dissociated cells were cultured in dishes coated with poly-D-lysine (Sigma) and containing Dulbecco’s modified Eagle’s medium (DMEM)–F12 (1:1, v/v) supplemented with B27 (Invitrogen) and recombinant human FGF2 (Invitrogen) at 20 ng/ml. For explant culture, the dissected telencephalon of ICR mouse embryos was cultured in DMEM-F12 supplemented with B27 and recombinant human FGF2. After culture of cells or explant tissue for 6 h, half of the medium was removed and replaced with medium supplemented with B27, human FGF2, and either Smoothened agonist (SIGMA-ALDRICH), recombinant human BMP4 (R&D Systems) or CHIR-99201 (Wako). Smoothened agonist was dissolved in dimethyl sulfoxide at a concentration of 5 mM and was added to culture medium at a final concentration of 2 μM. The recombinant BMP4 protein was dissolved at a concentration of 100 µg/ml in sterile 4 mM HCl containing 0.1% bovine serum albumin and was added to the culture medium at a final concentration of 50 ng/ml. CHIR-99201 was dissolved in dimethyl sulfoxide at a concentration of 1 mM and was added to culture medium at a final concentration of 5 µM. The cells or tissue were cultured for an additional 24 h and then frozen in liquid nitrogen before analysis.

### Statistical analysis

Data are presented as means ± s.e.m. and were compared with two-tailed Student’s paired or unpaired *t* test or by analysis of variance (ANOVA) followed by Bonferroni’s multiple-comparison test.

