## Supplemental Figures for "The Polycomb group protein Ring1 regulates dorsoventral patterning of the mouse telencephalon"

**Figure S1**

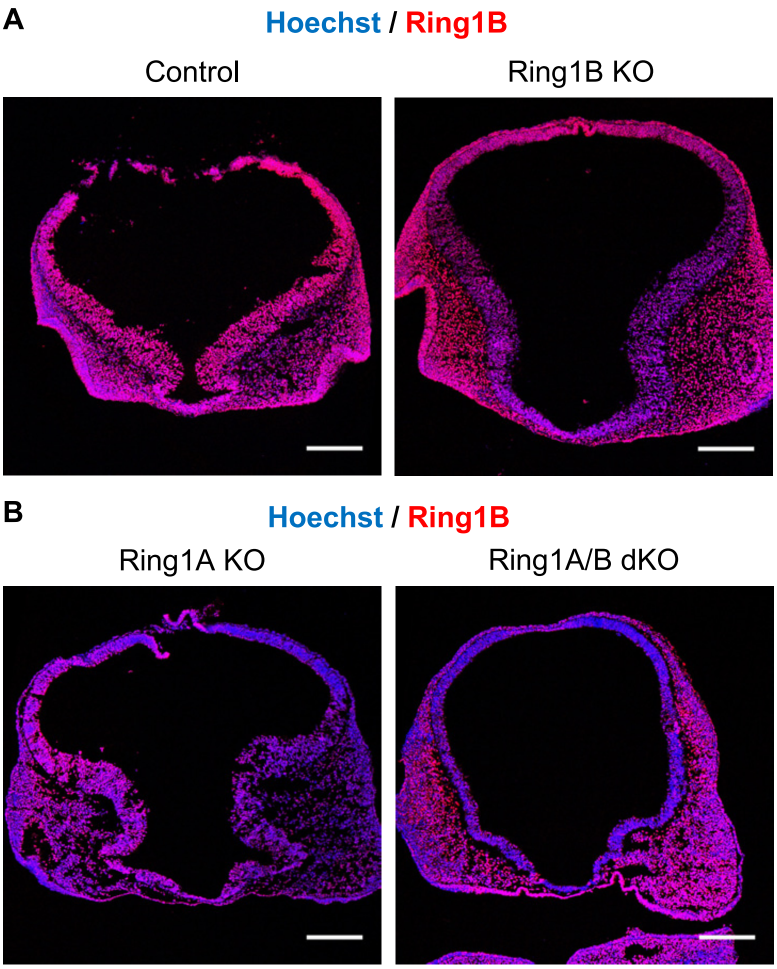


**Figure S1. Deletion of *Ring1* in neural tissues decreases the expression of *Ring1b* in the telencephalon**

(A, B) Coronal sections of the brain of control or Ring1B KO mice (A) or Ring1A KO or Ring1A/B dKO mice (B) at E10 were subjected to immunohistofluorescence staining with antibodies to Ring1B. Nuclei were counterstained with Hoechst 33342. Scale bars, 200 µm.

**Figure S2**

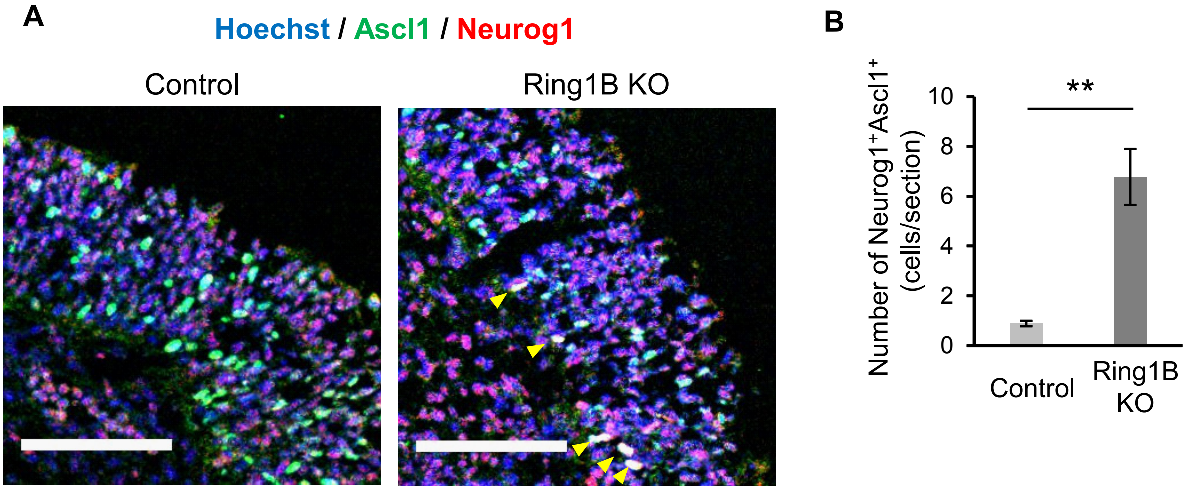


**Figure S2. Deletion of *Ring1b* increases Neurog1$^+$Ascl1$^+$ cells in telencephalon.**
(A) Higher magnification of boxed regions in Figure 4A (200 by 200 µm). Yellow arrowheads represent Neurog1$^+$Ascl1$^+$ cells. Scale bars, 80 µm.
(B) The average number of Neurog1$^+$Ascl1$^+$ cells in the control and Ring1B KO mouse telencephalon was determined with images similar to those in Figure 4A. Multiple sections of the telencephalon along the rostrocaudal axis were examined for each embryo. Data are means ± s.e.m. of values for three embryos. **P < 0.01 (two-tailed Student's unpaired *t* test)
