## Supplemental Table 2 for "The Polycomb group protein Ring1 regulates dorsoventral patterning of the mouse telencephalon"

**Supplementary Table 2, Primer sequences for qPCR and sequences for ISH probes**

**Primer sequences for RT-qPCR**

| **Gene** | **Forward primer** | **Reverse primer** |
| --- | --- | --- |
| *β-actin* | AATAGTCATTCCAAGTATCCATGAAA | GCGACCATCCTCCTCTTAG |
| *Nkx2.1* | GATGAGTCCAAAGCACACG | CTCCATGCCCACTTTCTTGTA |
| *Gsx2* | ACATACCTAAACCTGTCAGAGA | CTTGCAGCTTGTGTGATTG |
| *Emx1* | GTTCCCAGAGGCCATGA | TGGCCAAAGAAGCGATT |
| *Bmp4* | CTGAACTGAGTGCCATTTCC | CTCTACCACCATCTCCTGATAA |
| *Id1* | TACGACATGAACGGCTGCTA | TCTCCACCTTGCTCACTTT |
| *Wnt7b* | ACCAAAACTTGCTGGACCAC | ACGTGTTGCACTTGACGAAG |
| *Wnt8b* | CGTTCTTCTAGTCACTTGTGT | GGTCCCAAGCAAACTGGTATTTA |
| *Axin2* | GGTTCCGGCTATGTCTTT | CTCTCTCTGGAGCTGTT |
| *Ptch1* | TAAGAGTTTCAGCAATGTGAAGTATG | TGAAGCCAGTCTCTAAAGTAGT |
| *Gli1* | TGCCTGGAGAGACACAAT | TGGGCACCTCATGTAGC |
| *Shh* | GCCATCTCTGTGATGAACC | AGTGTAGAGACTCCTCTGAAT |

**Primer sequences for ChIP-qPCR**

| **Gene** | **Forward primer** | **Reverse primer** |
| --- | --- | --- |
| *β-actin* | CGGTTTGGACAAAGACCC | AAAGCCGTATTAGGTCCATC |
| *Gapdh* | TGCAGTCCGTATTTATAGGAACC | CTTGAGCTAGGACTGGATAAGCA |
| *Hoxa1* | CTGAACTGGCAAGAGGTGA | CGACCACGCAGAGATTT |
| *Hoxd3* | ACCTATTTGCGGTCGTC | CAAATTCGCCTGGGAAATCA |
| *Bmp4* | AGATACTCAGGCTGGGTT | AGAGAAGGAAGGAGTAGATGT |
| *Bmp7* | CAGCCTTCACCCAGAATG | CACCCTAGGACTTCAAGAG |
| *Wnt7b* | GGCATCCAGGAGTCAGA | CGCTATGGATGGAGCTAC |
| *Wnt8b* | ACTGTTTGGGATCGCTTAC | ACAAGTGACTAGAAGAACGC |

**Sequences for ISH probes**

*Shh*	NM_009170.3, 339-1652 (CDS 1-1314)

*Axin2*	NM_015732.3, 2069-2846 (CDS 1720-2497)
